# The beneficial role of *Candida intermedia* and *Saccharomyces boulardii* yeasts on the immune response of mice vaccinated with a SARS-CoV-2 experimental vaccine

**DOI:** 10.1101/2021.08.30.458196

**Authors:** Renan Eugênio Araujo Piraine, Neida Lucia Conrad, Vitória Sequeira Gonçalves, Jeferson Vidart Ramos, Fábio Pereira Leivas Leite

## Abstract

Non-*Saccharomyces* yeasts emerge as possible new probiotics with a beneficial effect equal to or greater than the reference probiotic yeast, *Saccharomyces boulardii*. In this work, we evaluated the immunomodulation effect caused by *Candida intermedia* in mice vaccinated with inactivated SARS-CoV-2. We conducted preliminary tests using murine macrophages (RAW 264.7) stimulated with viable and heat-killed yeast cells, culture supernatant, and DNA, using qPCR to detect the mRNA transcription. Next, mice were supplemented with *C. intermedia* before each dose of the SARS-CoV-2 vaccine, and then antibody production was measured by ELISA. The probiotic strain *S. boulardii* CNCM I-745 was used as a control. We also explored the differences in fecal microbiomes between the non-supplemented and supplemented groups. Live cells of *C. intermedia* increased the transcription of *IL-4*, *IL-13*, and *STAT3* by macrophages RAW 264.7, while heat-killed cells up-regulated *TNFα* and *Bcl6*, and the culture supernatant positively impacted *TLR2* transcription. Concanavalin, zymosan, and lipopolysaccharide were used to stimulate splenocytes from *C. intermedia*-supplemented animals, which showed increased transcription of *TNFα*, *IFNγ*, *IL-4*, *Bcl6*, and *STAT3*. Sera from these animals showed enhanced levels of anti-SARS-CoV-2 IgG, as well as IgG1 and IgM isotypes, and sIgA in fecal samples. The microbiome of the *C. intermedia*-supplemented group showed a higher abundance of *Bacteroides* spp. and *Clostridium* spp., impacting the Bacteroidetes/Firmicutes balance. We concluded that *C. intermedia* and *S. boulardii* could stimulate and impact the gene expression of cells important for innate immunity, influence the composition of the gastrointestinal microbiome, and primarily boost the humoral response after vaccination.

**Statements and Declarations Funding:** The present work was carried out with the support of Conselho Nacional de Desenvolvimento Científico (CNPq, Brazil), grant number 150538/2021-9.

## 1. Introduction

The definition of probiotics, which refers to live microorganisms that confer health benefits to the host when administered in adequate amounts [1], has been established for years. This definition was mainly based on the observation of the beneficial effect on the host that certain microorganisms cause in the prevention and treatment of pathological conditions [2]. However, knowledge of the influence on the general physiology of the organism, such as the modulation of immunological homeostasis, activation of mucosal immunity mechanisms, and modulation of adaptative immune functions, confers the aggregation of “immunobiotics” to the definition of these microorganisms [3].

Bacteria of different genera, such as *Lactobacillus* spp. and *Bifidobacterium* spp., comprise most of the probiotics characterized so far, however, the possibility of horizontal transmission of antibiotic resistance genes between probiotic and pathogenic bacteria makes their application limited [4, 5]. Yeasts are naturally antibiotic-resistant, but they are not capable of transmitting this resistance to other yeasts or bacteria because the genes involved in this role are present in their chromosomes, and not in bacterial plasmids, as occurs in tetracycline resistance genes, which can be shared between bacteria [6]. *Saccharomyces boulardii* has been commercialized and studied since the 1950s and is currently the best-characterized yeast in terms of its probiotic activity, safety, and application in the treatment of chronic gastrointestinal diseases and conditions [2]. Different strains of *S. cerevisiae* are also studied for their probiotic mechanisms of adhesion to intestinal cells, antagonism to bacteria, and immunomodulation [7], however, other non-*Saccharomyces* yeasts emerge as possible new probiotics with a beneficial effect equal to or greater than those observed for *Saccharomyces* yeasts [8]. Antagonism to pathogenic microorganisms, production of killer toxins, gastrointestinal tract (GIT) tolerance, and other characteristics that characterize probiotic microorganisms are also observed in yeasts of genera such as *Candida, Hanseniaspora, Pichia, Kluyveromyces, Issatchenkia*, *Torulaspora*, among others [9–11].

Microorganisms with probiotic functions can also impact the GIT microbiota. Researchers suggest that this impact is related to the effect on the composition and function of the intestinal microbiota, mainly by maintaining the stability of the microbial community, modulating metabolic functions, or favoring the presence of determined bacteria, fungi, and archaea taxa [12, 13]. Besides that, these probiotics positively impact intestinal surface integrity, producing antimicrobial compounds and metabolites that hinder pathogens’ growth and compete for binding sites in the intestinal mucosa [1]. The administration of probiotic yeasts has already been linked to positive changes in the microbiota of fish [14], pigs [15], cattle [16], and humans [1], among others. Since these changes in the microbiota composition can benefit and improve the immune response of hosts [1], this is another aspect that should be studied in the use of probiotics.

Recent studies developed by our research group with wild yeasts isolated from the environment in the region of Pelotas (Rio Grande do Sul, Brazil) [11], demonstrated the potential of probiotic activity of the yeasts *Hanseniaspora uvarum* (PIT001), *Pichia kluyveri* (LAR001) and *Candida intermedia* (ORQ001), as well as the ability to participate in fermentation processes of beverages. These non-*Saccharomyces* yeasts were able to tolerate conditions similar to the GIT *in vitro*, in addition to showing antimicrobial activity to foodborne pathogens such as *Salmonella* Typhimurium, *Listeria monocytogenes, Staphylococcus aureus, Pseudomonas aeruginosa,* among others. Strains of these three yeasts are generally associated with their presence in the microbiome, fermentative profile, and sensory contribution in fermented products such as wine, beer, and cocoa [17], and they currently arouse interest regarding their antagonism activity to other competitive or contaminant microorganisms [18–20].

Some strains of *C. intermedia* have demonstrated the ability to decrease or suppress the growth of *L. monocytogenes* [8, 21], in addition to being responsible for the production of antimicrobial peptides that affect other yeasts [20]. This yeast had its genome recently sequenced [22, 23], but its presence in fermented dairy foods was already described 30 years ago [24]. In a previous study [11], we identified *in vitro* some potential probiotic characteristics in the isolate *C. intermedia* ORQ001, such as high levels of auto-aggregation, co-aggregation with pathogenic Gram– and Gram+ bacteria, and a low decrease in cell viability after exposure to GIT conditions. Based on these results, we focused on the development of new studies regarding the probiotic potential of this yeast.

Although there are several studies evaluating the probiotic activity of specific strains, information regarding their administration and behavior *in vivo* is scarce, as well as knowledge of pathways and roles of action in GIT performance. The mechanisms of action of the immunomodulatory effect are still not completely elucidated, mainly for non-*Saccharomyces* yeasts, however, there is enough evidence that suggests the need for a deep investigation into it [25]. As highlighted for yeasts like *Pichia pastoris* [26] and by our group for *S. cerevisiae* and *S. boulardii* [27–29], the evaluation of immune system modulation and ability to control pathogens *in vivo* represent important and expressive results in the search for probiotic characteristics in these microorganisms. When these yeasts are administered as supplements, it is possible to observe the improvement of specific immune responses to vaccines [29], immunostimulatory activity [30], reduction of pathogen concentration in experimentally infected animals [28], or even the prevention of infections [27, 31].

Recently, researchers have linked the immunostimulatory effect of probiotics to the potentiation of the immune response to different vaccines and antigens, suggesting a great relevance in the treatment and prevention of infection by SARS-CoV-2, the virus responsible for the COVID-19 pandemic (Coronavirus disease) [32–34]. Previous studies developed by our research group using an experimental vaccine composed of inactivated SARS-CoV-2 [35] demonstrated that the vaccine stimulated humoral and cellular immune responses, which it is believed can be potentiated by the administration of probiotics. Thus, here we evaluated some potential probiotic characteristics through *in vitro* and *in vivo* tests, regarding the stimulation of macrophages by live cells of *C. intermedia* ORQ001 and *S. boulardii* CNCM I-745 and their derivatives (inactivated cells, culture supernatant, and yeast DNA), the immunomodulatory effect, and the impact on the gastrointestinal microbiome performed by oral supplementation with these yeasts, using as a target the immune response of mice vaccinated with inactivated SARS-CoV-2.

## 2. Material and methods

### 2.1. Strains and culture conditions

The wild isolate of *C. intermedia* ORQ001 was previously characterized regarding its probiotic potential by Piraine *et al.* [11], and for this study, it was obtained from the microorganism bank of the Microbiology Laboratory in the Federal University of Pelotas, as well as the commercial yeast *S. boulardii* CNCM I-745 (Floratil ®, a reference probiotic strain), which were cryopreserved in glycerol at −80 °C. Yeasts were grown overnight in YM (Yeast and Malt Extract) medium (0.3% yeast extract, 0.3% malt extract, 0.5% peptone, and 1% glucose) at 30 °C under constant agitation of 150 rpm. Successive steps of propagation under the previous conditions were conducted to scale up the yeast cultures, to a final concentration of 1 × 10^8^ CFU/ml (Colony-Forming Units). Cells were concentrated by centrifugation at 2.000 × g for 10 min using a DAIKI DTR-16000 centrifuge, counted by sequential serial dilutions, and stored at 4 °C until their use.

### 2.2. RAW 264.7 macrophage stimulation with *C. intermedia, S. boulardii,* and their derivatives

#### 2.2.1. Stimuli preparation

Live and inactivated (heat-killed) cells, culture supernatants, and DNA extracted from each yeast were used to stimulate the murine macrophage-like cell line RAW 264.7 (ATCC® TIB-71TM). Yeast cells were washed twice with Phosphate Buffered Saline (PBS) and then 10^8^ CFU/ml of viable cells were stored to be used subsequently during macrophage stimulus. Cell-free supernatant from yeast culture media was centrifuged and then separated in aliquots to further stimulate macrophages. Yeast cells at the same concentration (10^8^ CFU/ml) were inactivated by heat and pressure (heat-killed cells), autoclaving at 120 °C with a pressure of 1 atm for 20 min. After inactivation, samples were seeded onto YM agar medium and incubated for 48 h at 28 °C, a control step to assure correct yeast inactivation.

Total DNA from yeasts was extracted following an adaptation of the protocol described by Preiss *et al.* [36]. A volume of 3 ml from yeast cultures was centrifuged at 12.000 × g for 1 min, then it was suspended using 200 ul of breaking buffer solution (2% Triton-X 100 w/v, 1% SDS w/v, 100 mM NaCl, 100 mM Tris pH 8.0, EDTA 1 mM pH 8.0). An amount corresponding to 100 ul of glass microbeads (0.5 uM, Sigma-Aldrich) and 200 ul of phenol/chloroform/isoamyl alcohol (25:24:1) were added to the previous solution. Tubes were then vortexed for 2 min, and TE buffer (10 mM Tris pH 8.0, 1 mM EDTA pH 8.0) was added after that. Centrifugation was performed at 13.000 × g for 5 min, then the aqueous phase (∼350 ul) was transferred to a new tube, where 1 ml of ethanol 96% was added, homogenized, and incubated at room temperature for 10 min (without shaking). Tubes were centrifuged at 13.000 × g for 2 min, supernatants were discarded, and pellets were dried at room temperature for 20 min. DNA was eluted with 50 ul of DNAse free water, then DNA quality was evaluated by electrophoresis on 1% agarose gel (100V, 500 mA, 1 h), and its concentration was quantified by Nanovue^TM^ (Biochrom).

#### 2.2.2. RAW 264.7 cells culture and stimulation

Macrophages RAW 264.7 were grown as monolayers according to Santos *et al.* [37]. Briefly, cells were incubated in Dulbecco’s Modified Eagle Medium (DMEM) supplemented with 10% (v/v) Fetal Bovine Serum (FBS) at 37 °C in a 90% humidity atmosphere with 5% CO_2_ until approximately 80% confluence in the culture plate. Stimulation occurred in a yeast: RAW cells ratio of 10:1, following the adaptation of the protocol developed by Smith *et al.* [38]. RAW 264.7 cells were kept under stimulation for 24 h, with incubation in DMEM supplemented with 10% (v/v) FBS at 37 °C in an atmosphere of 90% humidity with 5% CO_2_. As a negative control, cells were stimulated with a DMEM medium only. As positive controls, Concanavalin A – ConcA (10 µg) and Zymosan (100 µg) (Sigma-Aldrich) were used. For the assay, 100 µl of live cells or 100 µl of heat-killed cells per well were used as stimuli, at a final concentration of 10^7^ CFU/ml. The same volume of supernatant (100 µl) was used, while for the DNA of each yeast, the concentration of 850 ng/well was targeted. The total volume of stimulus in the culture medium was 1 ml per well, carried out in triplicates.

### 2.3. Animal supplementation with live yeasts and vaccination with an experimental vaccine composed of inactivated SARS-CoV-2 virus

#### 2.3.1. Animals, supplementation, and vaccination

Ninety Balb/c mice of an age of 4–6 weeks were provided by the Central Animal Facility of the Federal University of Pelotas, and all procedures performed followed the guidelines of the Brazilian College of Animal Experimentation (COBEA) and were approved by the Ethics Committee on the Use of Animals at UFPel (CEEA n° 011015/2022-75). Mice were divided into nine experimental groups with ten animals each, as shown in Table 1.

**Table 1.** Experimental design of the groups of animals used in this experiment

| Group | Animals (n) | Supplementation | Vaccine |
| --- | --- | --- | --- |
| A | 10 | - | - |
| B | 10 | *C. intermedia* | - |
| C | 10 | *S. boulardii* | - |
| D | 10 | - | - |
| E | 10 | *C. intermedia* | - |
| F | 10 | *S. boulardii* | - |
| G | 10 | - | Inactivated SARS-CoV-2 virus |
| H | 10 | *C. intermedia* | Inactivated SARS-CoV-2 virus |
| I | 10 | *S. boulardii* | Inactivated SARS-CoV-2 virus |

| Group | Animals (*n*) | Supplementation | Vaccine |
| --- | --- | --- | --- |
| A | 10 | - | - |
| B | 10 | *C. intermedia* | - |
| C | 10 | *S. boulardii* | - |
| D | 10 | - | - |
| E | 10 | *C. intermedia* | - |
| F | 10 | *S. boulardii* | - |
| G | 10 | - | Inactivated SARS-CoV-2 virus |
| H | 10 | *C. intermedia* | Inactivated SARS-CoV-2 virus |
| I | 10 | *S. boulardii* | Inactivated SARS-CoV-2 virus |

Animal supplementation was performed once a day by oral administration (gavage) of 500 µl of yeasts *C. intermedia* or *S. boulardii,* at a concentration of 1 × 10^8^ CFU/ml. The supplementation was performed five days before each vaccine dose. For the non-supplemented group, the same volume (500 µl) of sterile PBS was administered to maintain equal stress conditions as the supplemented groups. Throughout the experiment, the animals were fed a commercial diet free of chemotherapeutics (Nuvilab® CR1 irradiated) and administered *ad libitum*.

The experimental vaccine was elaborated with formaldehyde-inactivated SARS-CoV-2 virus (kindly provided by Prof. Fernando Spilki, from Feevale University), at a concentration of 1 × 10^6^ PFU (Plate-forming Units), added with 10% aluminum hydroxide as an adjuvant. The mice were inoculated subcutaneously twice, with an interval of 21 days between vaccinations and a vaccine dose of 100 µl. For animals in which SARS-CoV-2 was not administered (unvaccinated groups), they were inoculated with a suspension composed of 100 µl of PBS with 10% aluminum hydroxide.

Blood samples were collected by the submandibular puncture on days 0, 7, 14, 21, 28, 35, and 42, with serum being separated by centrifugation at 5.000 × g for 5 min and then stored at – 20 °C until analysis. Fecal samples were collected and pooled (samples from all the animals in the group) before the first day of supplementation and following the same blood sample collection schedule, then stored at – 20 °C until their use. The schedule containing all the procedures and time points is presented in Fig. 1. Mice from groups A, B, and C were euthanized on day 0, while animals from the other groups were sacrificed on the 42^nd^ day (end of the experiment).

**Fig. 1.**
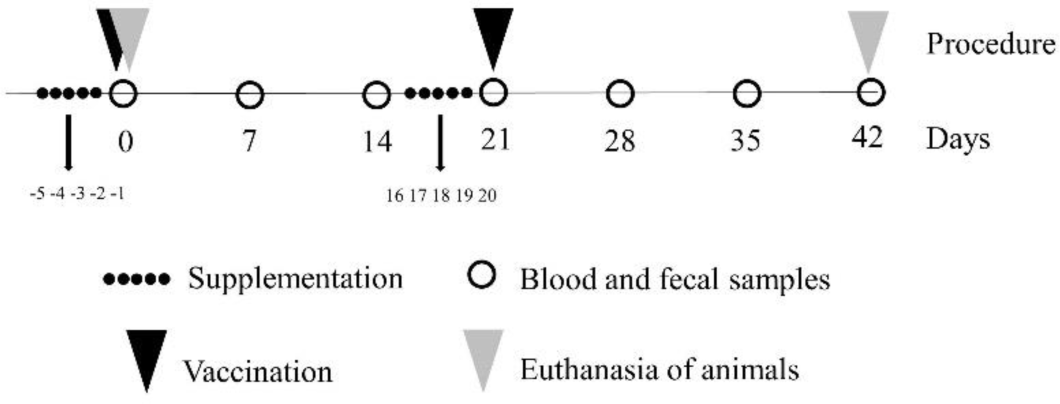
Schedule of animal experimentation. The experimental plan was conducted until day 42, consisting of a collection of blood and fecal samples every 7 days, supplementation with *C. intermedia* or *S. boulardii* by oral administration (days −5 to −1, and 16 to 20), and vaccination with two doses of inactivated SARS-CoV-2 virus (days 0 and 21)

#### 2.3.2. Cellular response to immune response stimulants with different origins

To evaluate the effect of yeast supplementation on the cellular immune response of the animals, the quantitative PCR method (qPCR) was used to amplify fragments of cytokine genes, transcription factors, and receptors in the cDNA obtained from mRNA. At day 0, just after the first five days of supplementation, spleens from mice in groups A, B, and C were pooled in duplicates of five each, to evaluate the immediate effect on the immune response after the period of administration of *C. intermedia* and *S. boulardii*.

The spleens were removed from animals and macerated, and then their cells (splenocytes) were suspended in a balanced HANK’S solution (without Ca^2+^ and Mg^2+^ ions). Cells were centrifuged and pellet suspended in lysis solutions (0.8% ammonium chloride), followed by a new step of wash and suspension in RPMI 1640 (Cultilab®) with 10% FBS (Cultilab®), totaling a standard concentration of 2 × 10^6^ cells/ml. These cells were cultivated in 24-well plates (Kasvi®), 1 ml per well, and incubated for 24 h at 37 °C with 5% CO_2_. After this period, the medium was renewed, and cells were stimulated in different ways: Concanavalin A (ConcA) (10 µg), Zymosan (100 µg), and Lipopolysaccharide (LPS) (10 µg). The stimuli were carried out during 18 h of incubation under the same conditions as before, then the supernatant was discarded, and cells were collected with the TRIzol® reagent (Sigma-Aldrich®) and stored at –70 °C. RNA extraction from splenocytes was performed according to the protocol provided by the manufacturer using the TRIzol method.

At the end of the experiment (42^nd^ day), duplicates of five spleens from mice from each of the other groups (D, E, F, G, H, and I) of vaccinated and non-vaccinated animals, supplemented or not, were used to obtain splenocytes, to evaluate cellular response patterns when stimulated with 1 x 10^5^ PFU/ml of inactivated SARS-CoV-2 virus. Incubation and stimulus conditions (time, temperature, culture medium), and RNA extraction were maintained as previously described.

The reaction for cDNA synthesis was performed using 400 ng of RNA, following the instructions available in the High-Capacity cDNA Reverse Transcription Kit (Applied Biosystems ®). Real-time Polymerase Chain Reaction (qPCR) was conducted in a Stragene Mx3005P real-time PCR system (Agilent Technologies®) to analyze the relative expression of the cytokine genes Interleukin 2 (*IL-2*), Tumor Necrosis Factor α (*TNFα*), Interferon (*IFNɣ*), Interleukin 4 (*IL-4*), Interleukin 12 (*IL-12*), Interleukin 13 (*IL-13*), Interleukin 23 (*IL-23*), Toll-like Receptor 2 (*TLR2*), and transcriptions factors *Bcl6, NFκβ,* and *STAT3*. The *β*-actin and GAPDH genes were used as endogenous reference controls. The sequence of cytokine-specific primers as well as qPCR conditions have been described previously [39, 40]. All samples were analyzed in triplicate. Relative expressions were calculated by comparing Threshold Cycle (Ct) values of *β*-actin and targeted genes with the non-supplemented group, according to the 2^-ΔΔCT^ method described by Livak and Schmittgen (2001).

#### 2.3.3. Analysis of humoral immune response

Indirect enzyme immunoassay (ELISA) was used to detect anti-SARS-CoV-2 antibodies in the sera of vaccinated mice. Briefly, microtitre plates (96-well, Cral®) were coated with inactivated SARS-CoV-2 virus (1 × 10^5^ PFU/ml) diluted in 0.1 M Carbonate-Bicarbonate buffer pH 9.8 overnight at 4 °C, washed three times with PBS-T (Phosphate-buffered saline + 0.05% Tween 20), and then incubated for 2 h at 37 °C with 100 µl/well of powdered milk 5% diluted in PBS. After a new step of PBS-T washing, pooled sera from each group were added in triplicate at a 1:100 dilution, for 2 h at 37 °C. Anti-IgG secondary antibodies conjugated with Horseradish Peroxidase (HRP) (Sigma-Aldrich®) at a dilution of 1:5.000 were applied after five PBS-T washes, and incubated again at the same conditions as previously described. The ELISA plates were washed again, and substrate buffer (0.4 mg ortho-phenylenediamine, 15 µl H_2_O_2_, and 0.1 M phosphate citrate buffer pH 4.0) was added with 100 µl/well to reveal the reaction. After 15 min in the dark at room temperature (∼25 °C), it was added 50 µl/well of 2 N sulphuric acid to stop the reaction. Absorbance was measured in a microplate reader (Thermoplate®) with a 492nm filter. Moreover, for antibody titration, using the same serum samples (D28, D35, and D42), two-fold dilutions were made in a range of 1:100 to 1:6400. The *cut-off* for antibodies titers we defined as the absorbance of day 0 plus the standard deviation.

The Mouse Monoclonal Antibody Isotyping Reagents Kit (Sigma-Aldrich®) was used for the evaluation of IgG1 and IgM isotypes, following the protocol suggested by the company. In this test, plates were coated with the SARS-CoV-2 virus as described before, and then serum samples of days 28, 35, and 42 from each group were pooled, diluted as described previously, and applied in triplicates. Secondary isotype-specific goat antibodies conjugated to HRP were used in 1:5.000 dilution, and plates were read as described before. For the detection of sIgA (secretory IgA isotype) in fecal samples, we conducted the ELISA test protocol described by Santos *et al.* [42]. For this purpose, 0.1 g of pooled fecal samples from days 28, 35, and 42 were suspended in 1% PBS with 1 mM Phenylmethylsulfonyl fluoride (PMSF, Sigma-Aldrich®) and 1% Bovine Serum Albumin (BSA), and mixed by vortex until complete homogenization. Thus, these samples were diluted with PBS-T + 5% powdered milk in a 1:2 ratio, then added (100 µl/well) over a plate coated with the SARS-CoV-2 virus. Specific sIgA antibodies to the SARS-CoV-2 virus were detected with goat anti-mouse IgA alpha-chain + HRP (Abcam®), diluted to 1:1.000. Incubation periods, washing steps, and solutions used for ELISA tests described before were also used for sIgA ELISA.

#### 2.3.4. Gastrointestinal microbiome evaluation

Total DNA present in feces from samples on day 0 (after 5 days of yeast supplementation) was isolated and sequenced using New Generation Sequencing by the Neoprospecta company (Brazil) to characterize the microbiome of the gastrointestinal tract. The amplicons were sequenced in paired-end mode (2×300bp) with Miseq Reagent Kit V3 R (600 cycles) on the Miseq Sequencing System platform (Ilumina®). Sequencing of the V3/V4 regions of the ribosomal RNA gene was performed with primers 341F (sequence CCTACGGGRSGCAGCAG), and 806R (sequence GGACTACHVGGGTWTCTAAT). Negative and positive controls were used during all processes. The raw data obtained from sequencing were submitted to quality control, taxonomic classification, visualization, and description of communities using the Knomics-Biota system [43].

Using the *R* programming language and *microbiome* package, the OTUs (*Operational Taxonomic Units*) table was normalized by “clr-transformation” to calculate alpha and beta diversities [44]. Alpha diversity was determined using Observed, Shannon, Simpson, InvSimpson, Fisher, and Evenness indices. The Kruskal-Wallis test was inferred to verify significant differences among groups (p<0.05). Beta diversity was estimated by Principal Component Analysis (PCA), using the Aitchison matrix [45].

#### 2.3.5. Statistical analysis

Serology data and those related to cellular response were analyzed by analysis of variance (2way ANOVA) with Dunnett’s multiple comparison test, and the statistical difference was determined if the *p-* value < 0.05. All analyzes were performed mainly in the statistical software GraphPad Prism 11 v. 7.

## 3. Results

### 3.1. Immunostimulatory activity of viable yeast cells and derivatives on RAW 264.7 cell culture

Live cells of *C. intermedia* were responsible for a significant stimulation (p<0.05) of the *IL-4* and *IL-13* mRNA production, with an expressive increase of 7.6 and 2.4-fold, respectively. On the other hand, down-regulation was observed for *IL-2* and *TNFα,* decreasing their expression (p<0.05) by 2 and 1.6-fold, respectively. Moreover, we observed that *TLR2* transcription was down-regulated (2.2-fold decrease) and, more expressively, the transcription factor STAT3 by 3.4-fold. Meanwhile, we found that viable cells of *S. boulardii* were responsible for an important stimulus, up-regulating the mRNA transcription of cytokines *IL-2* (8.6-fold), *IL-4* (3.0-fold), *IL-13* (8.7-fold), and *IL-23* (7.4-fold). Observing the expression profile for the other genes analyzed, there were no significant differences (p>0.05) in mRNA transcription after the stimulus with *S. boulardii* (Fig. 2).

**Fig. 2.**
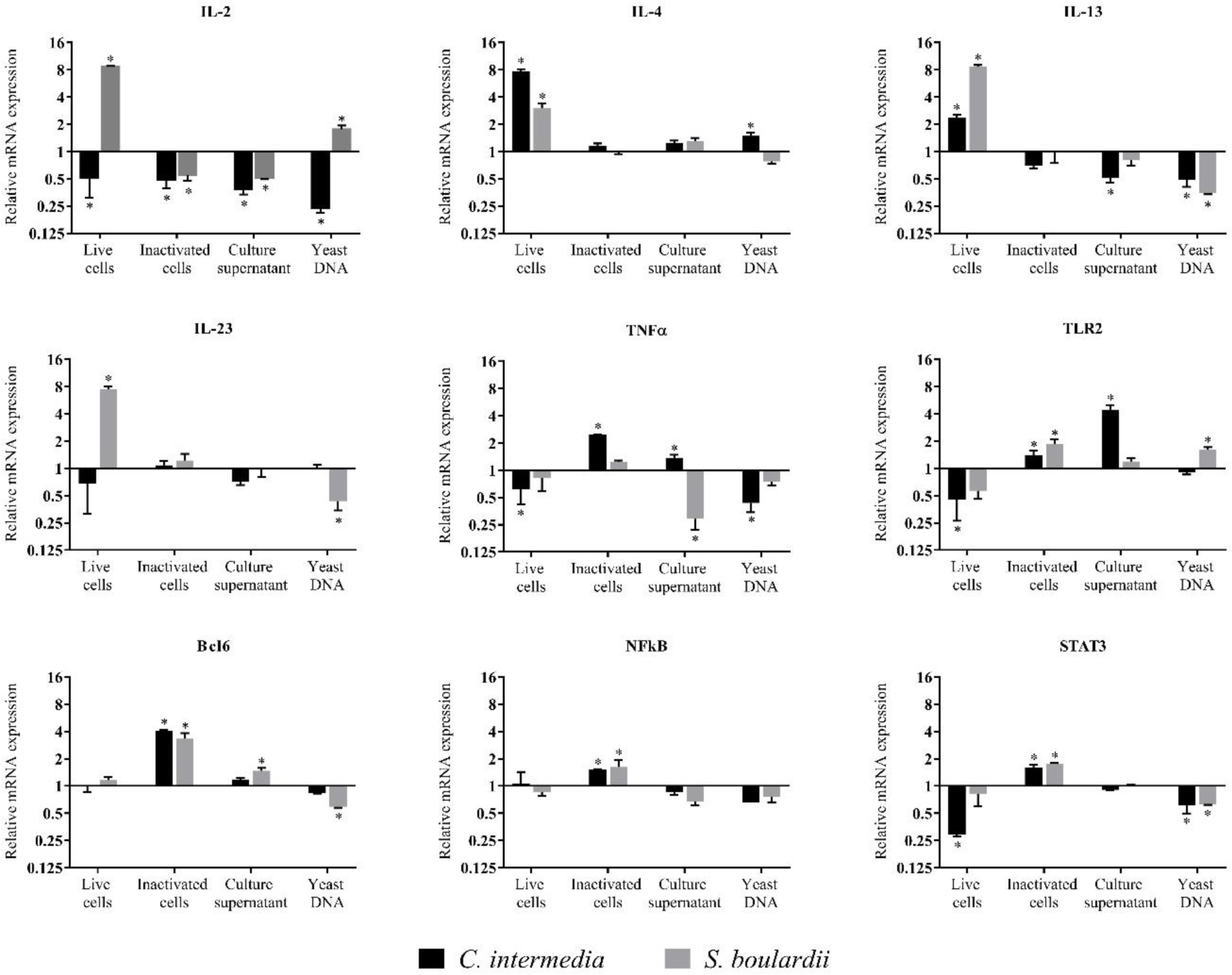
Gene transcription of cytokines, TLR2, and transcription factors during RAW 264.7 stimulation with yeast cells and their derivatives. Viable yeast cells, heat-killed inactivated cells, culture supernatant, and fungal DNA from *C. intermedia* ORQ001 and *S. boulardii* CNCM I-745 were used to stimulate RAW 264.7 macrophages. Relative mRNA transcription of *IL-2, IL-4, IL-13, IL-23, TNFα, TLR2, Bcl6, Nfκβ,* and *STAT3* was normalized using *β-actin* transcription level as reference. Data are shown as the mean ±SD (Standard Deviation). The symbol “*” means a significant difference (p<0.05) from the basal expression

After the stimuli with inactivated cells of *C. intermedia* and *S. boulardii*, a shift in the mRNA transcription profile was observed compared with the stimuli with viable yeast cells. In this case, killed cells of *C. intermedia* were not able to stimulate an increase in *IL-4* and *IL-13* transcription, but an up-regulation was identified for *TNFα* transcription (2.4-fold increase). Differently from what was observed for live cells, the stimulus with inactivated cells induced a potentiation in the mRNA transcription of *Bcl6* (4.0-fold increase) and, in a more discrete way (but still significant), *NFκβ* (1.5-fold), *STAT3* (1.6-fold), and TLR2 (1.4-fold). Inactivated cells of *S. boulardii* also caused a down-regulation of IL-2 (1.8-fold decrease), which contrasts with that observed when live cells of this yeast were used as a stimulus for RAW 264.7 macrophages. Both cell receptor and transcription factors transcriptions were enhanced with inactivated *S. boulardii* cells, with increases of 1.8 times for *TLR2*, and between 3.3 and 1.6 times for *Bcl6*, *NFκβ,* and *STAT3*.

The supernatant of *C. intermedia* culture was also used to evaluate the response generated by *in vitro* cultured macrophages. We observed that *C. intermedia* culture supernatant induced an increase in *TLR2* transcription (a 4.4-fold increase) and a decrease in *IL-2* and *IL-13* levels (a 2.6 and 2.0-fold decrease, respectively). The supernatant of *S. boulardii* culture also revealed an impact on the mRNA transcription of macrophages RAW 264.7, down-regulating the transcription of cytokines *IL-2* (2.0-fold) and *TNFα* (3.4-fold), and up-regulating Bcl6 transcription factor (1.5-fold). Finally, we found that the DNA extracted from *C. intermedia* discreetly stimulated an increase in IL-4 production (1.5-fold), but mainly the inhibition of *IL-2* by a 4.1-fold decrease in mRNA transcription, with less intensity for *IL-13*, *TNFα*, and *STAT3* between 1.6 to 2.3-fold. Likewise, *S. boulardii* DNA has also been shown to impact the mRNA transcription of some genes, positively stimulating *IL-2* (1.7-fold) and TLR2 (1.6-fold), and negatively stimulating IL-13 (2.9-fold), IL-23 (2.3-fold), Bcl6 and STAT3 (1.6-fold both).

### 3.2. Modulation of immune cells response by a short period of supplementation with yeasts

After five days of supplementation with *C. intermedia* and *S. boulardii*, splenocytes from animals from each group were collected and cultured *in vitro*. These cells were stimulated with ConcA, Zymosan, and LPS, with mRNA transcription of cytokines (*TNFα, IFNγ, IL-4, IL-12, IL-23*), transcription factors (*Bcl6, NFκβ, STAT3*), and *TLR2* being evaluated by qPCR (Fig. 3). We identified that splenocytes from animals supplemented with *C. intermedia* showed increased mRNA transcription of *TNFα, IFNγ,* and *IL-4* for all stimuli when compared to non-supplemented animals, in which the biggest difference levels were observed for *TNFα* in ConcA-stimulated cells (6.7-fold increase) and *IFNγ* in LPS-stimulated cells (20.4-fold increase). For some gene transcription profiles, the mRNA transcription could even result in a change from down-regulation to up-regulation when animals were supplemented with *C. intermedia*, as was observed for *IL-4* cytokine mRNA transcription in LPS-stimulated cells. For *IL-23* cytokine transcription, it was also detected an increase in its level in splenocytes from animals supplemented with the non-*Saccharomyces* yeast.

**Fig. 3.**
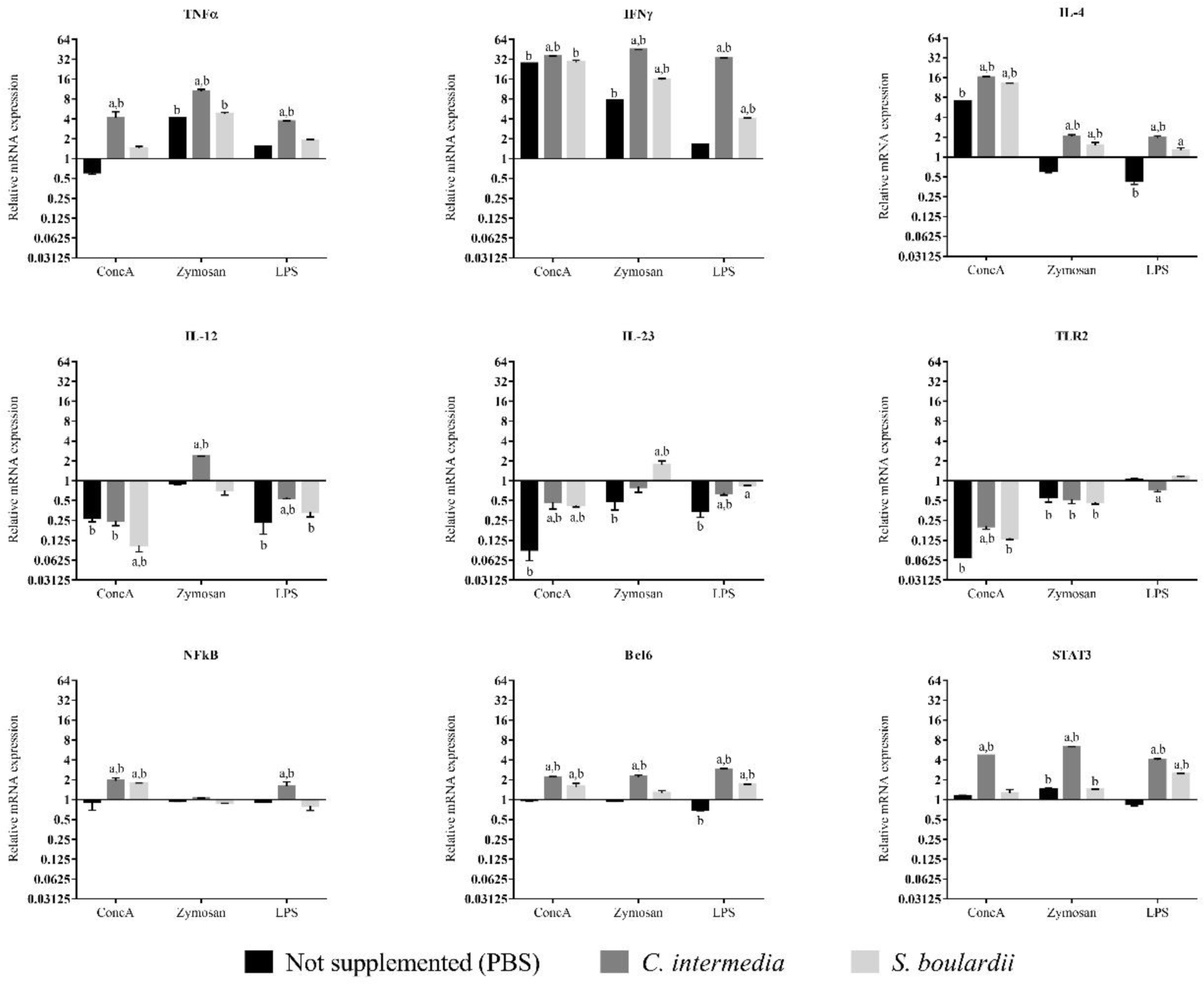
Gene transcription of cytokines, TLR2, and transcription factors during splenocyte stimulation with ConcA, Zymosan, and LPS. Splenocytes were obtained from animals supplemented after a short schedule (five days) with *C. intermedia* and *S. boulardii,* or non-supplemented (control). Relative mRNA transcription of *TNFα, IFNγ, IL-4, IL-12, IL-23, TLR2, Bcl6, Nfκβ, and STAT3* was normalized using *β-Actin* transcription level as a reference. Data are shown as the mean ±SD (Standard Deviation). The letter “**a**” means a significant difference (p<0.05) from the non-supplemented group, and “**b”** represents a significant difference (p<0.05) from the basal expression level

The stimulation of splenocytes from the *S. boulardii* group generated a response of cytokine transcription similar to that observed for the *C. intermedia* group; however, it was detected in some cases at lower levels than the non-*Saccharomyces* group or with no statistical difference for the non-supplemented group. The only two cases in which a different behavior was identified were on *IL-12* mRNA transcription (splenocytes from the *S. boulardii* group presented a 10-fold decrease in the cytokine level) and *IL-23* (an up-regulation with a 1.7-fold increase). While there was no statistical difference for *TLR2* transcription in stimulated cells from animals supplemented with *S. boulardii* and non-supplemented, cells obtained from *C. intermedia-*supplemented animals demonstrated a level below the basal transcription in ConcA stimulus, however, it was not as low as observed for non-supplemented animals (a 5.0 and a 14.2-fold decrease, respectively).

The mRNA transcription of transcription factors was also altered. *Bcl6* and *STAT3* transcription was up-regulated after ConcA, Zymosan, and LPS stimuli in cells of animals supplemented with *C. intermedia*, with an increase ranging between 2.2 to 4.6-fold. An up-regulation was also observed for *NFκβ* after ConcA and LPS stimuli; however, it was more discrete (between 1.6 and 1.9-fold increase) than that observed for the other transcription factors. Supplementation with *S. boulardii* impacted *NFκβ* and *Bcl6* in the ConcA stimulus (∼1.6-fold increase), as well as in *Bcl6* and *STAT3* transcription levels in LPS-treated cells (1.7 and 2.4-fold increase, respectively).

### 3.3. Immunomodulation of humoral and cellular response of animals vaccinated with SARS-CoV-2 inactivated virus

Observing the production of antibodies specific to the SARS-CoV-2 virus (total IgG), the *C. intermedia-*supplemented group showed greater antibody levels (p<0.05) than the non-supplemented group after 14 days of the first vaccine dose, with similar levels (p>0.05) on the 21^st^ day (day of the second vaccine dose). The animals supplemented with *S. boulardii* also presented greater IgG SARS-CoV-2-specific levels than the non-supplemented group, however, on day 28 only, with no statistical difference (p>0.05) between their absorbance values for the rest of the time points. The most interesting results were found for the *C. intermedia*-supplemented group starting at day 28, in which higher absorbance values were detected, representing an enhanced antibody level that lasted until day 42. Compared to the non-supplemented group on days 28, 35, and 42, sera from animals that ingested *C. intermedia* during two rounds of supplementation showed an increase of 41, 28, and 22% (respectively) in the absorbance values of anti-SARS-CoV-2 IgG at these time points (Fig. 4).

**Fig. 4.**
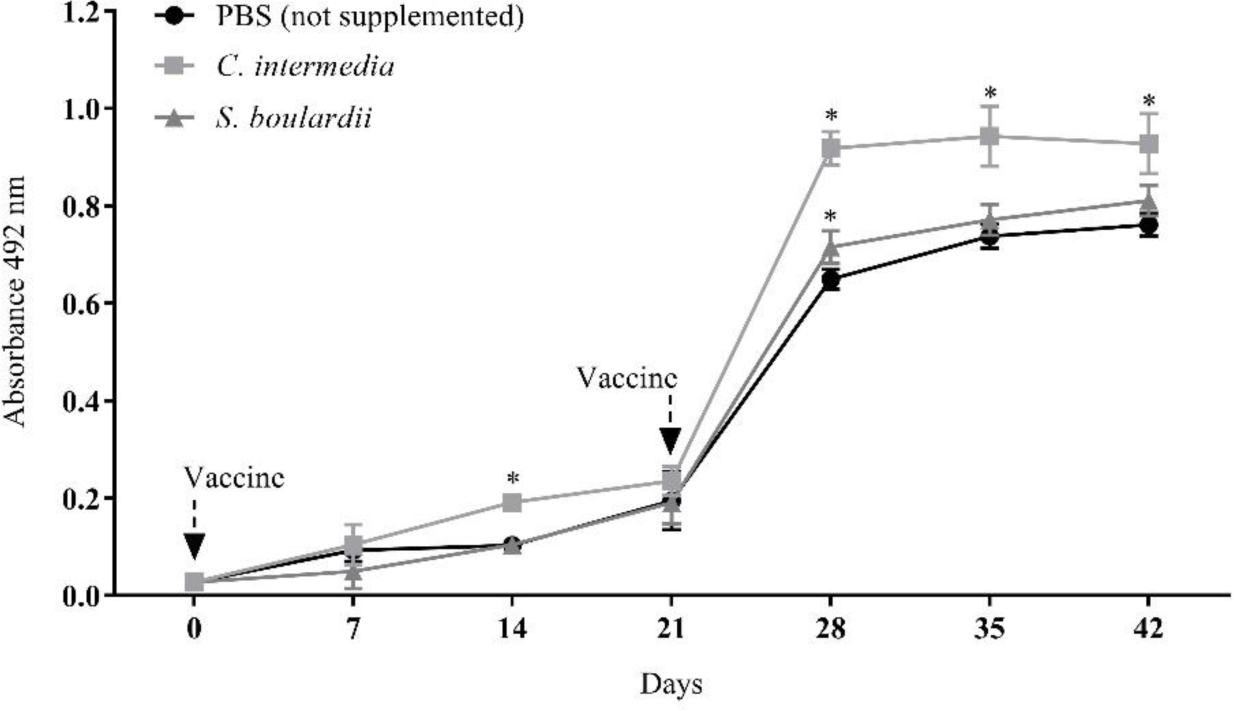
Humoral immune response (total IgG) of the animals non-supplemented (control) and supplemented with *C. intermedia* and *S. boulardii*, and vaccinated with inactivated SARS-CoV-2. The data represent the means (± standard deviation) of the absorbance values (492_nm_) obtained by indirect ELISA, using pooled sera collected every seven experimental days. The statistical analysis was performed by 2-way ANOVA, and if there was a statistical difference (p<0.05) from the non-supplemented group (PBS), an indication with an “*” is presented In the IgG titration (Fig. 5), for sera collected on the 28^th^ day, the antibody titer for the non-*Saccharomyces* group (3.200) was 2 times higher than that observed in the non-treated (non-supplemented) group (1.600). On day 35, the *C. intermedia*-supplemented group had an antibody titer of 3.200, which was four times higher than the non-supplemented animals (800). On the last experimental day (day 42), sera from animals supplemented with *C. intermedia* were the only ones that maintained the titer of 3.200, while the non-supplemented group repeated the previous titer of day 35 (1.600). Sera from animals supplemented with *S. boulardii* demonstrated the same titers as those identified in the non-supplemented group for days 28 and 35, however, a decrease was detected on day 42 for the *S. boulardii* group, in which its titer was determined as 800.

**Fig. 5.**
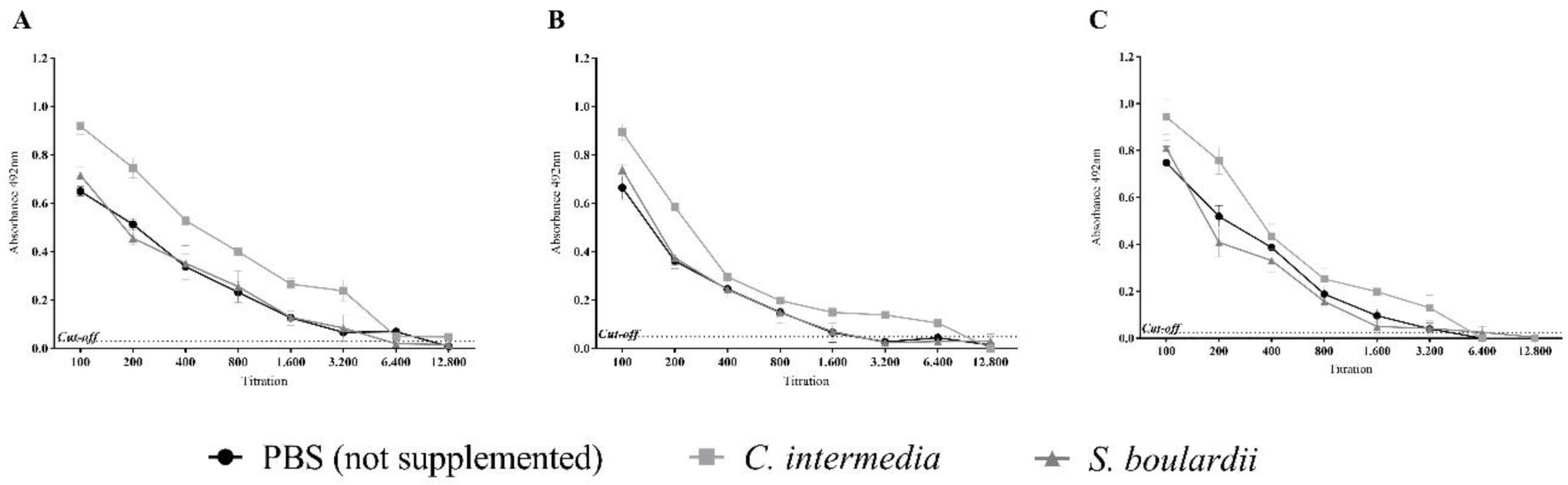
Titration of total IgG levels in mice vaccinated with inactivated SARS-CoV-2 and previously supplemented with *C. intermedia* and *S. boulardii* yeasts. A) Titration of serum samples collected on experimental day 28. B) Titration of samples from the 35^th^ day. C) Titration of samples from the 42^nd^ day. The data represent the means ± standard deviation of the absorbance values obtained by indirect ELISA. The titer was considered the highest pooled sera dilution detected above the *cut-off* (p<0.05) Possible modulation of the humoral response mediated by the yeasts was also monitored by the evaluation of antibody isotypes. Analyzing the IgG1 isotype (Fig. 6A), it was observed for day 28 that all groups presented the same levels in pooled sera, with no statistical difference (p>0.05) from the non-supplemented group. However, for samples collected on the 35^th^ day, it was detected at superior levels (p<0.05) of IgG1 in *C. intermedia-*supplemented animals and in the *S. boulardii*-supplemented group, which presented lower levels of this isotype. On the 42^nd^ day, sera from non-supplemented and *C. intermedia-*supplemented animals showed similar levels of IgG1, while those from *S. boulardii*-supplemented animals remained below these levels.

Levels of IgM isotype were also evaluated in the pooled sera of vaccinated mice on days 28, 35, and 42. No statistical difference was observed among groups on the 28^th^ experimental day; however, on days 35 and 42, *C. intermedia*-supplemented animals demonstrated higher levels of IgM in their sera than animals from non-supplemented and *S. boulardii* groups (Fig. 6B). It was possible to detect sIgA presence in the fecal samples from treated animals, an indication that there was seroconversion and stimulation of a specific mucosal response to the SARS-CoV-2 vaccine. The non-supplemented group demonstrated the highest levels of sIgA at day 28, while there was a gradual increase in sIgA levels for the *C. intermedia* group during the analyzed time points, reaching the maximum value (comparing all groups) on the 42^nd^ day. Inversely, *S. boulardii* fecal samples demonstrated the lowest levels of sIgA since the 28^th^ experimental day (Fig. 6C).

**Fig. 6.**
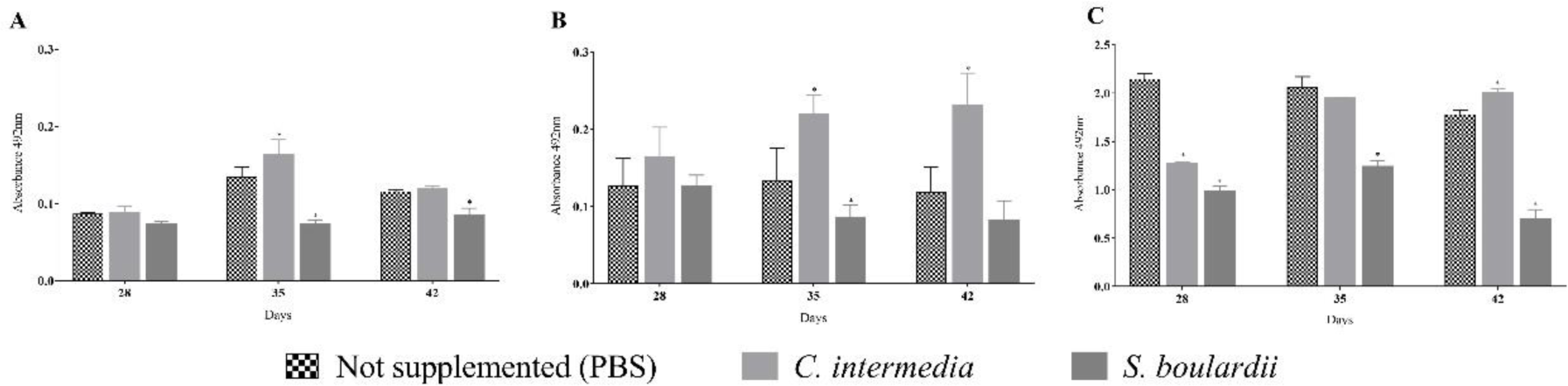
Isotypes IgG1, IgM, and sIgA specific to SARS-CoV-2 produced by vaccinated mice, supplemented and non-supplemented with yeasts. A) IgG1 detection on pooled sera of animals in the 28^th^, 35^th^, and 42^nd^ experimental days. B) IgM detection on pooled sera of animals in the 28^th^, 35^th^, and 42^nd^ experimental days. C) sIgA detection on fecal samples of animals in the 28^th^, 35^th^, and 42^nd^ experimental days. The data represent the means ± standard deviation of the absorbance values obtained by indirect ELISA. The statistical analysis was performed by 2-way ANOVA, and if there was a statistical difference (p<0.05) from the non-supplemented group (PBS), an indication with an “*” is presented

**Fig. 7.**
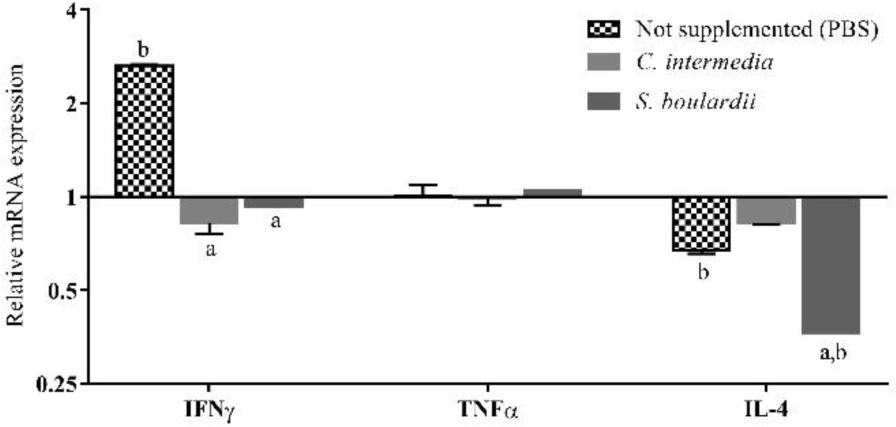
Relative mRNA transcription of *IFNγ, TNFα*, *and IL-4* cytokines after the stimulation of splenocytes with inactivated SARS-CoV-2. Splenocytes from non-supplemented (control group), *C. intermedia*-supplemented, and *S. boulardii-*supplemented animals were cultured *in vitro*, stimulated with SARS-CoV-2, and evaluated for the mRNA transcription of pro-inflammatory (*IFNγ* and *TNFα*) and anti-inflammatory (*IL-4*) cytokines. The relative mRNA transcription of these cytokines was normalized using the *β-actin* transcription level as a reference. Data are shown as the mean ±SD (Standard Deviation). The letter “**a**” means a significant difference (p<0.05) from the non-supplemented group, and “**b**” represents a significant difference (p<0.05) from the basal expression level

The cellular response in the vaccinated animals (supplemented or not with yeasts) was also evaluated by the stimulus of *in vitro*-cultured splenocytes with inactivated SARS-CoV-2. It was observed that the non-supplemented group (control) presented the highest transcription levels for *IFNγ* (p<0.05) when compared to the other groups, showing a 2.6-fold increase in basal mRNA transcription after the SARS-CoV-2 stimuli (Fig. 7). Meanwhile, splenocytes from *C. intermedia* and *S. boulardii* groups showed no statistical difference in the transcription levels for *IFNγ* mRNA from the basal level. Both treated and control groups showed similar mRNA transcription levels of the other pro-inflammatory cytokine, *TNFα*, with no statistical difference (p>0.05) from basal levels in its transcription. The mRNA transcription of the anti-inflammatory cytokine *IL-4* by the stimulated splenocytes was detected for the *C. intermedia-* supplemented and non-supplemented groups as similar (p>0.05) or close to basal levels (p<0.05) (1.5-fold decrease), respectively. However, a more pronounced impact on its transcription was detected in splenocytes from *S. boulardii*-supplemented animals, which demonstrated a 2.7-fold decrease in *IL-4* transcription.

### 3.4. Gastrointestinal tract microbiomes of supplemented and non-supplemented animals

The microbiome of fecal samples from treated and non-treated groups revealed some important differences in bacterial composition, which were suggested to be related to supplementation with *C. intermedia* and *S. boulardii*. We observed in the control group (non-supplemented, with a normal diet) that the microbial environment on GTI is composed, at the phylum level, of Bacteroidetes (53.9%) and Firmicutes (44.8%), with less abundance (<1% each) for Actinobacteria and Proteobacteria. Analyzing the genera found, *Bacteroides* spp. (42.0%), *Lactobacillus* spp. (35.9%) and *Parabacteroides* spp. (7.9%) (Fig. 8A) were the most abundant among the bacterial OTUs classified.

**Fig. 8.**
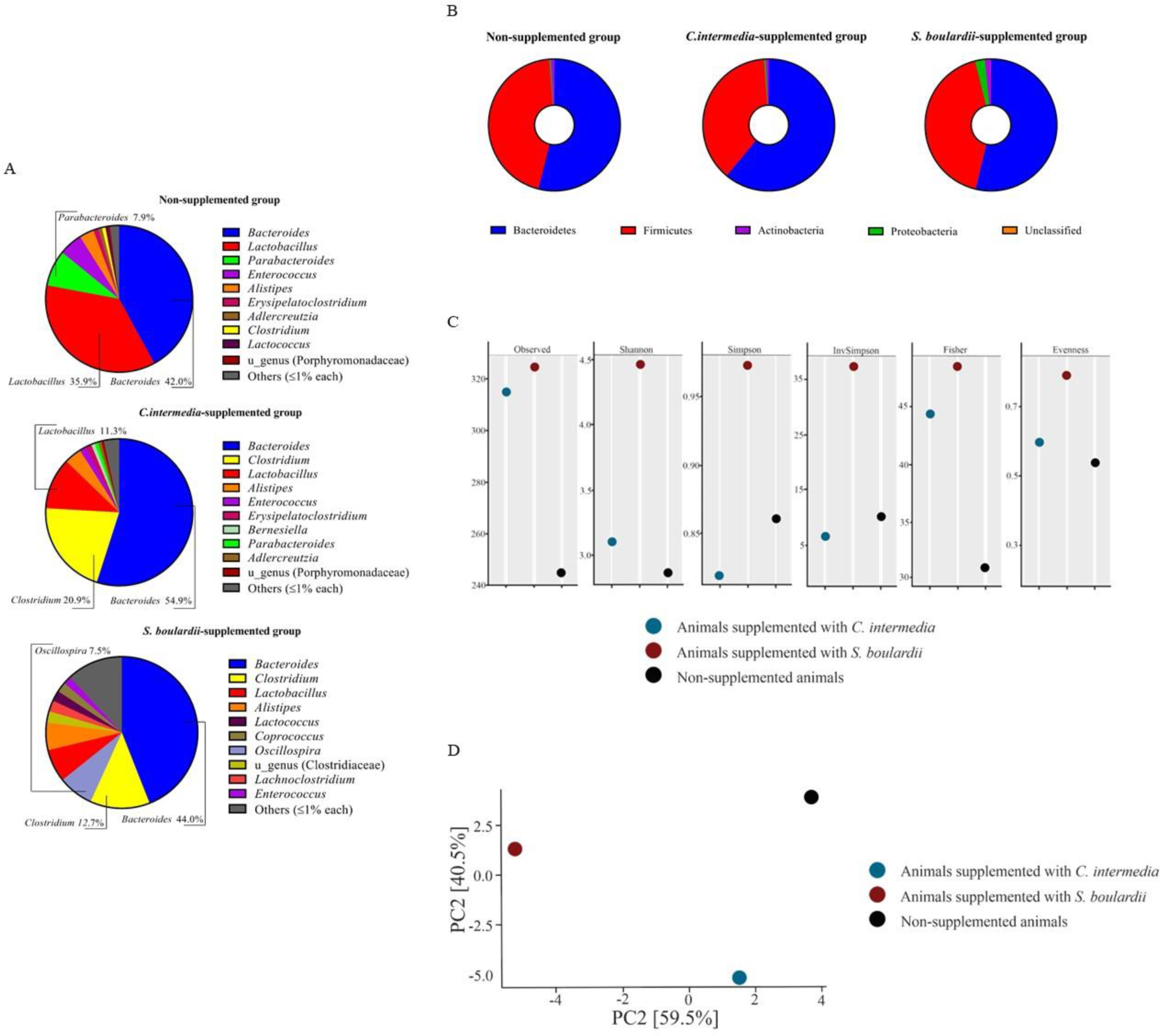
Microbiome analysis of the fecal samples from non-supplemented, *C. intermedia*-supplemented, and. *S. boulardii-*supplemented animals. After a 5-day schedule of supplementation, fecal samples were collected from different animals in each experimental group, pooled, and submitted for microbiome analysis. A) The most abundant bacterial genera in the sequenced samples, emphasizing the top 3 in each sample. B) Bacterial abundance at the phylum level. C) Alpha-diversity of the microbiomes, demonstrated by Observed, Shannon, Simpson, InvSimpson, Fisher, and Evenness indices, D) Beta-diversity, estimated by Principal Component Analysis (PCA)

It was detected in the microbiome of the *C. intermedia*-supplemented group a similar profile for the 10 most abundant genera compared to the non-supplemented group, however, we observed a higher abundance for *Bacteroides* spp. (54.9%) and a shift in *Clostridium* spp. and *Lactobacillus* spp. OTUs abundance, which was 20.9% and 11.3%, respectively. Thus, this observation impacted the Bacteroidetes/Firmicutes balance, showing an abundance of 61% of Bacteroidetes and 37.8% of Firmicutes (Fig. 8B). Meanwhile, the microbiome identified in the GTI sample obtained from *S. boulardii*-supplemented animals also showed a higher abundance of *Clostridium* spp. (12.7%), a similar level of *Bacteroides* spp. (44.0%), and a more pronounced presence of other genera with relevant abundance (≥1%) for the top 10 detected, such as *Oscillospira* spp. (7.5%). Although other genera were detected, the abundance of Bacteroidetes (53.7%) and Firmicutes (42.4%) was maintained in a similar proportion to that observed for the control group.

The alpha diversity of the microbiomes was measured by different indices. The species richness was detected through the observation of a total of 244 species being identified in the non-supplemented group sample, while samples of *C. intermedia-*supplemented animals showed 315 and *S. boulardii*-supplemented animals showed 325 species detected (Fig. 8C). While the non-supplemented group and the *C. intermedia*-supplemented group demonstrated in their samples close values for Shannon, Simpson, InvSimpson, and Evenness indices, the *S. boulardii*-supplemented group showed the highest values for these indices, as well as for Fisher (in which treated groups demonstrated closer values). In the beta-diversity analysis (Fig. 8D), we observed that the microbiome of animals supplemented with *C. intermedia* was less altered when compared to non-supplemented animals, while this alteration was more significant for the group supplemented with *S. boulardii*.

## 4. Discussion

Yeasts, especially those belonging to *Saccharomyces* spp., can participate in the activation of immune cells [38, 46, 47]. When macrophages (and other immune cells) are exposed to fungi, this interaction initiates intracellular signaling cascades that culminate in opsonization, phagocytosis, and the production of cytokines, chemokines, and antimicrobial peptides, among others [48]. The stimulation of macrophages by viable and non-viable cells depends on the cell structure of each yeast, cellular portions, yeast surface, internal cell components, and actively secreted molecules (by live cells) to the extracellular environment [46]. In addition, the medium supernatant (cell-free), obtained after yeast culture, also may have metabolic byproducts that interact with immune system cells [49].

This interaction still needs to be explored, as researchers have characterized different response patterns mediated by probiotics, pathogenic yeasts, and their components [50, 51]. Analysis of cytokine expression by human DCs (dendritic cells) by Bazan *et al.* [52] verified a response based on IL-12, IL-23, and IL-27 cytokines when stimulated with different yeast genera, including *Saccharomyces* spp. and *Candida* spp. In our study, viable cells of *S. boulardii* were able to stimulate macrophages to produce high transcription levels of *IL-2, IL-4, IL-13,* and *IL-23*. IL-2 promotes the growth and development of peripheral immune cells, initiating a defensive immune response through the survival and division of regulatory T cells (Treg) and proliferation of cytotoxic T cells [53]. Cytokines IL-4 and IL-13 play a fundamental role in the immunosuppressive and anti-inflammatory activity, while IL-23 has an immunoregulatory and pro-inflammatory role, sustaining cell-mediated responses focusing on intracellular infection elimination [53–55]. As noted by Stier and Bischoff [47], *S. boulardii* leads to a general unspecific immune system activation, and variations in patterns of cytokines stimulated can be observed according to cell lines analyzed (dendritic cells, macrophages, lymphocytes, etc.), stimulus, test conditions (*in vitro* × *in vivo*), and *in vivo* challenge with pathogens, among others.

A macrophage response based on *IL-4* and *IL-13* transcription was also observed for live *C. intermedia* cells, with a transcription level for *IL-4* 2.5-fold higher than that observed for *S. boulardii*. While *S. boulardii* is the most studied yeast for its immunostimulatory activity, new studies that relate this characteristic to non-*Saccharomyces* yeasts emerge with important findings [38]. Variations in metabolic activity and cell wall composition of *Candida* spp. lead to differences in phagocytosis and levels of cytokines being produced by immune cell lines stimulated *in vitro* [56]. There are extensive variations in yeast cell walls when comparing different fungal species and strains, such as α-glucans in addition to β-glucans, and differences in concentration of chitosan, galactomannans, and melanin [57]. Although cell wall composition between *C. intermedia* and *S. boulardii* is comparable, Lozančić *et al.* [58] demonstrated significant differences in their genera regarding patterns of GPI-anchored and non-covalently attached proteins, cell wall thickness, permeability, and amounts of mannan and glucans. Thus, different responses induced in immune cells may be related to cell wall complexity and its components [48].

Although Smith *et al.* [46] did not find differences in cytokine-inducing properties among live, UV-irradiated, and heat-killed cells, in our study, a considerable variation (p<0.05) in relative mRNA transcription was detected between stimuli with viable and non-viable cells. After a heat treatment associated with high pressure, cell inactivation occurs through membrane damage, loss of nutrients and ions, protein denaturation, and essential enzyme inactivation, which can lead to modifications in cell coarseness and roughness [59, 60]. Previous studies reported even higher expression levels of cytokines when macrophages were stimulated with heat-killed probiotics than viable cells, as observed by Miyazawa *et al*. [61]. In our study, the level of *TNFα* mRNA transcription was potentiated when macrophages were stimulated by heat-killed cells of *C. intermedia*, a behavior also observed for other *Candida* species by Navarro-arias *et al.* [56], who stimulated peripheral blood mononuclear cells (PBMCs) with viable and heat-killed *C. albicans, C. tropicalis, C. guilliermondii, C. krusei,* and *C. auris*.

Since Bcl6 plays a fundamental role in the regulation of Th2-type inflammation and is constantly expressed in monocytes [62], we sought to observe its transcriptional levels in stimulated macrophages. Bcl6 also regulates macrophage function by repressing pro-inflammatory cytokine production and controlling the differentiation of Th17 cells [63]. During active inflammation occasioned by infections, microorganisms can bind to PRRs (Pattern Recognition Receptors) and activate signaling pathways via MAP kinases and NFκβ, resulting in the stimulation of cytokines with pro-inflammatory activity [64]. Increased *NFκβ* transcription is involved with TLR2 receptor activity, which has its expression level increased by mannose and β-glycans recognition [65]. Toll-like receptors are innate immune response infection sensors that participate in the activation or inhibition of macrophage activity via the Jak-STAT pathway; signaling via STAT3 is activated by several cytokines and their receptors, such as IL-2, IL-6, IL-10, IL-23, and IL-27 [66]. The main role of STAT3 in macrophages is to mediate anti-inflammatory effects, restricting gene transcription of pro-inflammatory cytokines [67] and repressively impacting NFκβ signaling pathways [68]. Because these transcription factors are directly involved in signaling and activation pathways of the immune response, we have chosen them to better understand the impact of yeast recognition by immune cells. In our study, heat-killed cells were responsible for stimulating the highest levels of transcription of *Bcl6, STAT3, and NFκβ* mRNA in macrophages RAW 264.7.

Macrophages are usually inactive but can be activated through various stimuli during an immune response [51, 69–72]. Yeast cells and their cell wall components are generally important stimulators of TLR2 and Dectin-1 [73–75]. Low TLR2 expression from other stimuli does not necessarily impact cytokine expression, as Smith *et al.* [38] noted, since recognition of cell wall components may also depend on other receptors, such as Dectin-1 and mannose receptors. It could also be observed in our data, as although stimuli from *S. boulardii* and its derivatives resulted in low levels of *TLR2* transcription, cytokines like *IL-13* were highly transcribed when live *S. boulardii* cells were used to stimulate macrophage responses. Variations in TLR2 and cytokine expression levels occasioned by live and inactivated cells stimuli may be explained by β-glycans exposure on the entire yeast cell surface occurring after heat treatment, while intact cells usually expose β-glycans only through budding scars [51].

Molecules present in *C. intermedia* culture supernatants were also responsible for stimulating significant levels of TLR2, suggesting that metabolites secreted by yeasts are also important in stimulating receptors of immune system cells. Secreted proteins by *Candida* spp. yeasts are linked with TLR2/TLR4 recognition, as demonstrated by Wang *et al.* [76], promoting an inflammatory response in DCs and macrophages stimulated *in vitro*. The most common ligands related to TLR2 are PAMPs (Pathogen-Associated Molecular Patterns) originating from glycolipids, lipopeptides, or GPI-anchored structures [77], thus it is suggested that higher receptor mRNA transcription in these cases is related to secretome products or proteins detached from yeast cell walls that have these structures in their conformations [76, 78].

Probiotic microorganisms act in multiple ways, including the activation and stimulation of immune cells (lymphocytes, granulocytes, macrophages, mast cells, epithelial cells, and dendritic cells) to release various cytokines, regulating the innate and adaptive immune responses, and potentiating anti-inflammatory or pro-inflammatory responses, or even maintaining homeostasis [79]. Santos *et al.* [80] observed in PBMCs from animals supplemented with *S. boulardii* CNCM I-745 an increased mRNA transcription of *IL-2, IFNγ, and Bcl6* when stimulated by ConcA or *Clostridium chauvoei*. In our study, macrophages from animals supplemented with *S. boulardii* also showed an increase in *IFNγ* (after Zymosan and LPS stimuli) and *Bcl6* (ConcA and LPS) levels. Chou *et al.* [81] also demonstrated an up-regulation of IFNγ expression in spleen cells from animals supplemented with an *S. cerevisiae* fermentation product, showing that live cells plus their active secreted metabolites are important for the immunomodulatory effects of the *Saccharomyces* yeast.

Zymosan, ConcA, and LPS are well-known immune response stimulants with different origins [75, 82], so comparing splenocyte responses after stimulation with these molecules permits the prediction of and helps identify the influence of yeast administration on animals’ immune systems in different scenarios. Based on our findings, *C. intermedia* was suggested as a microorganism with potent immunomodulation effects in cytokine transcription by splenocytes, even superior to *S. boulardii*, which was observed in *TNFα, IFNγ, IL-4,* and *IL-12* mRNA transcription. IFNγ is one of the most potent macrophage activators, and together with TNFα and IL-12, it is a pro-inflammatory cytokine that promotes cell-mediated immunity [69]. Besides inducing an up-regulation in cytokine transcription, it was also observed an increase in the mRNA levels of transcription factors, of which STAT3 transcription was consistently up-regulated no matter what the stimulus. Since our work was the first to explore the immunomodulatory activity of *C. intermedia* and its application *in vivo*, more data is necessary to better understand all the immunological signals and pathways involved after its administration to animals.

When dealing with the *Candida* genus, it is always important to highlight that safety is crucial to determining the possibility of yeast usage as a tool to promote health, since even non-*albicans* species may cause infections [83]. Most of the infections (63-70%) related to *Candida* spp. are caused by *C. albicans*, and the rest are associated with other 18-30 species classified in *Candida* genus (non-*albicans*) that comprises around 200 species, such as *C. glabrata, C. tropicalis, C. parapsilosis, C. krusei, C. auris* [83, 84]. At this date, *C. intermedia* is not a common human pathogen; it is described in the literature only in 3 cases associated with infections involving immunosuppressed individuals [85, 86]. Nevertheless, a presumption of safety for use of *C. intermedia* in fermented food products was published in the Bulletin of the International Dairy Federation 495/2018 [87], and, in our study, there were no related deaths or apparent signals of yeast infection in the supplemented animals. Therefore, further studies are needed to confirm the safety of supplementation with *C. intermedia*.

Modulation of vaccine response by *C. intermedia* and *S. boulardii* ingestion was evaluated in our study associated with an experimental vaccine composed of inactivated SARS-CoV-2. The potential of *S. boulardii* to improve immune responses after vaccination was identified by our group in previous studies [29, 80], and is also related in the literature [88–90]. Our results demonstrated for the first time the possibility of supplementation with yeasts to improve the immune response to a SARS-CoV-2 vaccine, as well as showed non-*Saccharomyces* yeasts as potential (and promising) candidates for this application. While studies linking bacteria and their probiotic activity are more common, an unknown number of non-conventional yeasts (besides *Saccharomyces* spp.) are waiting to be discovered in this field [2]. *C. intermedia* supplementation suggested a positive impact on the humoral response of mice vaccinated with the anti-SARS-CoV-2 vaccine, an aspect that confirmed the immunomodulatory role of this yeast. Although our results do not reflect confirmation of higher protection against virus infection or organism protection, they provoke interest in new studies using other vaccines (viral, bacterial, recombinant, etc.), animal models (ovine, bovine, humans, etc.), pathogen challenge, and protocols of supplementation (short-term × long-term) associated with the yeast administration.

Some authors described enhanced levels of IgG and sIgA when probiotics were administered in association with immunizations using DNA vectors, recombinant subunits, and viral vaccines [80, 88, 91], which was also observed in our results for *C. intermedia*-supplemented animals. These findings are relevant because secretory IgA is predominant in mucosal surfaces, playing an important role in viral immunity, and it is suggested its role in an enhanced capacity to neutralize SARS-CoV-2 [92]. Klingler *et al.* [93] also described the importance of IgA antibodies; moreover, it was found that IgG1 and IgM have a strong contribution to SARS-CoV-2 neutralizing activity. After immunization with a SARS-CoV-2 vaccine, Ruggiero *et al*. [94] identified different IgG/IgM response patterns in vaccinated individuals, in which a coordinated response based on these two immunoglobulin classes was determined to be important to confer protection against viral infection. Here we observed that supplementation with *C. intermedia* increased the production of both IgG and IgM levels, a possible immune response modulation that may be a determinant to prevent or hinder viral infection.

Besides the influence on humoral and cellular responses, we investigated changes in the GIT microbiome caused by *C. intermedia* and *S. boulardii* supplementation. The intestinal microbiome, composed of several microorganisms, plays an essential role in host immunity and makes great contributions to the host’s health, including improving nutrient digestion and absorption, differentiation of the intestinal epithelium, maintenance of intestinal mucosal barrier, prevention of pathogen attachment, and impacts on the polarization of the intestinal-specific immune response [1, 15]. Supplementation of animals’ diets with *S. boulardii* is being studied mainly for farm animals as an alternative to antibiotics, in which its activity of shaping the gut microbiome is highlighted [15]. This change in microbiome composition caused by *S. boulardii* also impacts the decrease of obesity, fat mass, hepatic disorders, and metabolic inflammation, as demonstrated by Everard *et al.* [95]. While Everard *et al.* [95] and Yu *et al.* [96] described in their results a decrease in Firmicutes and an increase in Bacteroidetes abundance in mice supplemented with this yeast, we observed a similar Firmicutes/Bacteroidetes ratio between the non-supplemented and *S. boulardii*-supplemented groups.

On the other hand, differences in the Firmicutes/Bacteroidetes ratio were observed in the microbiome of *C. intermedia* animals, with increased levels of Bacteroidetes and lowered levels of Firmicutes. The increase in Bacteroidetes abundance, especially for the *Bacteroides* genus, may be related to the utilization of mannan available on *Candida* cell walls through degrading enzymes (mannanases and mannosidases) expressed by *Bacteroides* spp., serving as a nutrient (carbon) source for the bacteria [97]. LPS derived from *Bacteroides* spp. are described as showing impaired or even inhibitory capacity to elicit an inflammatory response, and a decrease in its abundance in the gut microbiome often results in an augmented population of pro-inflammatory bacteria [98, 99]. The enriched presence of certain Bacteroidetes (*Prevotella* spp. and *Bacteroides* spp.) with anti-inflammatory properties was linked to fewer adverse effects after vaccination and associated with the highest antibody titers in some cases [100], correlating with our results. However, this positive association is not a consensus in the literature.

Bacterial species can take advantage of the biofilm created by *Candida* and adhere to it to thrive under conditions that would be harsh for them in GIT [97, 101]. It was demonstrated for *Clostridium perfringens* and *C. difficile* that these bacteria benefit from the microenvironment and microbial interactions when co-cultured with *Candida albicans* [101], and in our study, this positive interaction between *Candida-Clostridium* may be the answer to enhanced levels of *Clostridium* spp. in the microbiome of *C. intermedia*-supplemented animals. The abundance of *Lactobacillus* spp. was lower in the microbiome of yeast-supplemented animals, which may be due to the competition between yeasts and bacteria for the same metabolic niches throughout the gastrointestinal tract [84]; however, further studies are needed to prove this linkage. Finally, several mechanisms affect the response to vaccines, including vaccine formulation, dose, immunization route, vaccination schedule, the host immune system, and the gut microbiota [90]. The modulation of resident microbiota by probiotics is a factor that may improve vaccine immunogenicity [90, 102], thus we suggest that the alterations detected in the microbiomes of the supplemented mice could be one of the key factors associated with the increase in the humoral response detected in these groups.

In summary, we demonstrated in this study that supplementation with the yeasts *C. intermedia* ORQ001 and *S. boulardii* CNCM I-745 could stimulate immune cells important in innate immunity, were able to regulate the expression of different genes related to immune response, influenced the GIT microbiome, and primarily boosted a specific humoral response after vaccination. Even though *S. boulardii* is a well-known yeast with probiotic activity, every new study is important to understand its immunomodulation effect. For the first time, *C. intermedia* was proposed as a microorganism that modulates the immune system, and it demonstrated a positive impact on the immune response of animals vaccinated with inactivated SARS-CoV-2. Our findings obtained through *in vivo* tests complement previous *in vitro* studies that already proposed its probiotic activity, provoking interest in further studies to investigate its safety. We also concluded that the administration of *Saccharomyces* and non-*Saccharomyces* yeasts may have importance in vaccination schedules, representing an alternative for boosting immune responses.

## Statements and Declarations

### Conflicts of interest

On behalf of all authors, the corresponding author states that there is no conflict of interest.

### Availability of data and material

All data generated or analyzed during this study are included in this published article.

### Author’s contributions

All authors contributed to the study’s conception and design. Material preparation, data collection, and analysis were performed by Renan E. A. Piraine, Neida L. Conrad, and Vitória S. Gonçalves. Renan E. A. Piraine and Jeferson V. Ramos conducted bioinformatics. The first draft of the manuscript was written by Renan E. A. Piraine and all authors commented on previous versions of the manuscript. All authors read and approved the final manuscript.

### Ethics approval

All procedures performed followed the guidelines of the Brazilian College of Animal Experimentation (COBEA) and were approved by the Ethics Committee on the Use of Animals at UFPel (CEEA n° 011015/2022-75).

### Consent to participate

Not applicable

### Consent for publication

Not applicable

## Acknowledgment

We would like to thank Prof. Dr. Fernando Spilki and his team for providing inactivated SARS-CoV-2 samples. We are grateful to all students and workers for their support during the work at the animal facility of UFPel.

## Notes

### Competing Interest Statement

The authors have declared no competing interest.

### Summary of Updates

The immunostimulatory effect of the yeasts was studied deeper through new tests; the previous results were re-evaluated and presented in a new way that improves their interpretation. For this purpose, we focused on the non-Saccharomyces yeast Candida intermedia and added results about its possible probiotic activity. We mainly explored the immunomodulatory effects of C. intermedia and S. boulardii. To achieve the new findings shown in this version, we administered these two yeasts to mice, which were vaccinated with a vaccine composed of inactivated SARS-CoV-2. Now we demonstrate the modulation of the cellular and humoral responses by C. intermedia and S. boulardii, amplifying the knowledge about non-Saccharomyces yeasts and suggesting that these yeasts may be as beneficial as the reference probiotic yeast.

